# Mild traumatic brain injury is associated with increased thalamic subregion volume in the subacute period following injury

**DOI:** 10.1101/2024.06.03.597265

**Authors:** Maggie E Baird, Richard Beare, Marc L Seal, Joseph Yuan-Mou Yang, Jacqueline F. I. Anderson

**Affiliations:** Melbourne School of Psychological Sciences, University of Melbourne, Victoria, Australia; National Centre for Healthy Ageing and Peninsula Clinical School, Monash University, Melbourne, Australia; Developmental Imaging, Clinical Sciences, Murdoch Children’s Research Institute, Parkville, Victoria, Australia; Department of Pediatrics, University of Melbourne, Victoria, Australia; Department of Neurosurgery, Neuroscience Advanced Clinical Imaging Service (NACIS), Royal Children’s Hospital, Parkville, Victoria, Australia; Neuroscience Research, Clinical Sciences, Murdoch Children’s Research Institute, Parkville, Victoria, Australia; Psychology Department, The Alfred Hospital, Prahran, Victoria, Australia

**Keywords:** mild traumatic brain injury, thalamus, thalamic subnuclei, volumetrics, diffusion weighted imaging

## Abstract

Structural vulnerability of the thalamus remains under investigated in mild traumatic brain injury (mTBI), and few studies have addressed its constituent nuclei using robust segmentation methods. This study aimed to investigate thalamic subnuclei volume in the subacute period following mTBI. Trauma control (TC) and mTBI patients aged 18 – 60 years old completed an MRI neuroimaging protocol including both high resolution structural (T1w) and diffusion weighted sequences at 6 – 11 weeks following injury (mean: 57 days; sd 11). Each thalamus was segmented into its constituent subnuclei, which were grouped into eight lateralised subregions. Volumes of the subregions were calculated. Neurite Orientation Dispersion and Density (NODDI) maps with parameters optimised for grey matter were computed for the same subregions. Group differences in subregion volumes and NODDI parameters were investigated using Bayesian linear modelling, with age, sex, and intracranial volume included as covariates. Comparisons of mTBI (n = 39) and TC (n = 28) groups revealed evidence of relatively increased grey matter volume in the mTBI group for the bilateral medial and right intralaminar subregions (BF_10_ > 3). Of the subregions which showed volume differences, there was no evidence for differences in NODDI metrics between groups. This study demonstrates that in the subacute period following mTBI, there is evidence of increased volume in specific thalamic subregions. Putative mechanisms underpinning the increased volume observed here are disordered remyelination, or myelin debris yet to be cleared.

**Significance statement:** Despite the prevalence of continued cognitive, somatic, and vestibular symptoms in the subacute period (6 – 12 weeks) following an mTBI, a clear neuropathophysiological profile is yet to be determined. One key vulnerable structure in mTBI could be the thalamus, a subcortical grey matter structure which comprises numerous subregions. The present study investigated whether changes in thalamic subregion volume are evident in the subacute period. For the first time, we show that at approximately 8 weeks following injury, mTBI is associated with increased volume in specific thalamus subregions. This provides an important avenue for continued investigation into the clinical significance of these findings.

---

Mild traumatic brain injury (mTBI) constitutes the majority of individuals who sustain a traumatic brain injury, with incidence estimates of at least 300/100,000 people per year (Cassidy et al., 2004). Despite a positive prognosis following the resolution of the acute neurological disruption typical of mTBI, incomplete recovery is common, with many patients continuing to report a heterogenous array of cognitive, affective, and vestibular symptoms in the months to years following injury (Machamer et al., 2022). The existence of divergent recovery trajectories in the mTBI population has driven interest in the factors underpinning poor recovery, which are currently limited by an incomplete understanding of injury-related neuropathology. In particular, a clear pathophysiological profile of mTBI in the subacute period (i.e., between 6 – 12 weeks) following injury is yet to be established. Considering the prevalence of symptom reporting beyond this period, it is important to clarify the nature and extent of structural change in the post-acute period. To date, neuroimaging research within the mTBI population has largely focused on quantifying the degree of white matter microstructural pathology following injury, with far less attention paid to subcortical structures such as the thalamus.

The thalamus is a bilateral grey matter diencephalic structure which forms the upper and lateral walls of the third ventricle. Although historical perspectives have conceptualised it as a relay station through which sensory information reaches the cortex, the thalamus is now understood to serve an important role in influencing subcortical and cortical dynamics by integrating information processing through its dense and widespread connectivity profile (George & Das, 2021). Indeed, virtually all cortical areas receive thalamic input and project back to the thalamus, thereby enabling its contributions to key cognitive, sensory, vestibular, and motoric functions (Antonucci et al., 2021).

Several lines of evidence point to the thalamus as a vulnerable structure in mTBI. Given its position at the core of the brain, the thalamus appears to be anatomically vulnerable to the rotational forces of mTBI in which the cerebrum rotates around the brain’s centre of mass (Shaw, 2002). During these rotational forces, the greatest tissue strain occurs in the midbrain and diencephalic structures, including the thalamus (Viano et al., 2005; Zhang et al., 2004). Animal histopathological evidence suggests the thalamus is a region of prominent inflammation and neurodegeneration following mTBI, and prolonged thalamic inflammation is evident up to 17-years following human traumatic brain injury (Dikranian et al., 2008; Johnson et al., 2013; Natale et al., 2002; Ramlackhansingh et al., 2011). Taking a symptom-informed rationale, given thalamic contribution to a range of functions, it is reasonable to consider that its vulnerability in mTBI could contribute to the many and varied mTBI symptoms.

Thalamic morphological features can be investigated *in vivo* using MRI-based techniques that first isolate the thalamus from its surrounding structures and then quantify macrostructural properties, such as volume. This has proven to be a useful technique to identify thalamic volume changes associated with a range of psychiatric and neurological disorders (Bora et al., 2012; Minagar et al., 2013). However, despite evidence suggesting its possible vulnerability in mTBI, the thalamus remains an under investigated structure in this cohort, and the findings of existing volumetric studies have been mixed. In the immediate acute period following injury, no evidence of thalamic volume differences between mTBI and healthy or orthopedic control groups has been reported, an unsurprising finding considering work suggesting volume changes unfold over a protracted timeline as secondary injury mechanisms occur (Cole et al., 2018; Hellstrom et al., 2017; Woodrow et al). In the subacute period, some studies report reduced thalamic volume at one- to two-months, (Zagorchev et al., 2016; Zhou et al., 2021), while another reported greater thalamic volume at six-months compared to healthy controls (Munivenkatappa et al., 2016). The longitudinal components of these studies support the notion that volume change is progressive over time following mTBI, although there is no clear consensus among previous research as to the temporal evolution and expected direction of change (Munivenkatappa et al., 2016; Zagorchev et al., 2016).

One key limitation of previous thalamic volumetric studies is the treatment of the thalamus as a uniform structure. While there is commonality in cellular properties across the entire thalamus, it is structurally heterogenous, with at least 50 subnuclei identified depending on the degree and method of classification (Tregidgo et al., 2023). These subnuclei contribute variably to function, enabled by differences in connectivity targets, neurochemical receptors and transporters, and aspects of cytoarchitecture (Antonucci et al., 2021; Keun et al., 2021). Previous work shows selective vulnerability of subnuclei across various conditions, such as multiple sclerosis, Parkinson’s disease, and schizophrenia, which indicates certain cellular responses may predominate in specific subregions (Bhome et al., 2022; Magon et al., 2020; Mørch-Johnsen et al., 2023). Given the heterogenous contributions of thalamic subnuclei to various functions, differential subnuclei sensitivity in mTBI may have implications for the functional deficits observed following injury. Therefore, there is strong motivation to address the thalamus in terms of its constituent subnuclei, to provide a more precise view of thalamic vulnerability in mTBI.

The trend to regard the thalamus as a unitary structure in MRI-based neuroimaging studies is underpinned in-part due to difficulty attaining meaningful subnuclei segmentations. This is because T1w images typically lack sufficient tissue contrast that can be exploited to define nuclei boundaries within the thalamus (Iglehart et al., 2020). Recent methodological advancements can overcome this limitation by employing joint T1 and diffusion MRI segmentation methods, which leverage complementary information from diffusion MRI data to improve boundary placement. A recently developed, well validated, unsupervised segmentation method presents a promising means to robustly segment the thalamus and its constituent subnuclei (Tregigdo et al., 2023). To date, this technique is yet to be applied in an mTBI sample.

Changes in grey matter volume alone do not provide biologically specific information as to the underlying cellular mechanisms and tissue microstructural properties. Therefore, it may be informative to supplement volumetric analyses with findings from an independent diffusion weighted imaging (DWI) method that can probe the arrangement and structure of dendrites and axons to allow for the inference of microstructural features. Among these diffusion models is Neurite Orientation Dispersion and Density (NODDI) imaging, which considers contributions to the DWI signal from distinct intraceullar, extraceullar and cerebrospinal fluid compartments, each of which has unique diffusion properties (Zhang et al., 2012). The NODDI model yields three metrics: neurite density index (NDI), a marker of dendrite and axonal density; orientation dispersion index (ODI), a metric of the degree of bending and fanning of neurites (i.e., their spatial organisation); and the volume fraction of isotropic diffusion compartment (ISO), which estimates the free water content (Zhang et al., 2012). While their interpretation is not unambiguous, NODDI metrics can allow for the inference of cellular responses to injury, and as such could provide complementary quantitative evidence for the mechanisms underpinning volume change (Kamiya et al., 2020).

Preliminary evidence suggests the thalamus may be a vulnerable structure in mTBI, but this is yet to be thoroughly investigated by methodically assessing structural change in subnuclei. The aim of the present study was to investigate thalamic nuclei macro- and micro-structural features in the subacute period following mTBI. To address this aim, a multimodal neuroimaging approach was applied to segment the thalamus into its constituent subnuclei. Subsequently, a complementary analysis of volume and NODDI parameters of the identified subregions was performed, from which structural neuropathology can be inferred.

## Materials and Methods

### Participants

Two groups of adults who were admitted due to traumatic injury, mTBI (n = 40) and trauma controls (TCs; n = 30), were recruited as inpatients between 2017 and 2023 from the Royal Melbourne and The Alfred hospitals in Melbourne, Australia. Participants were classified into the mTBI group if they fulfilled the World Health Organisation criteria for definition of mTBI: Glasgow Coma Scale score between 13 and 15 at 30 minutes post-injury, and one or more of the following symptoms: < 24 hours post-traumatic amnesia; < 30 minutes loss of consciousness, impaired mental state at time of accidence (e.g., confusion, disorientation); and/or transient neurological deficit (e.g., seizure). Participants were included in the TC group if their traumatic injury occurred in the absence of a head strike. All participants fulfilled the following inclusion criteria: ages 18 – 60, no active psychiatric episode at time of injury and no psychiatric treatment and/or symptoms for preceding 12 months, no previous diagnosis of psychotic disorder, no current or historical heavy alcohol use (average > 6 standard drinks/day), no current or historical intravenous drug use, no class A drug use in previous 12 months, and conversational English fluency. Injuries that occurred in the context of assault and self-harm were excluded. Veterans and professional athletes were excluded.

Additionally, mTBI participants were excluded if they had previously sustained more than two mTBIs, or ever sustained a traumatic brain injury of moderate-severe severity, and TC participants were excluded if they had ever sustained a traumatic brain injury of any severity. All participants provided informed consent. Demographic information including years of education and biological sex was also recorded. This study was approved by The Alfred and Royal Melbourne Hospital Human Research Ethics Committees.

## MRI data acquisition

MRI data was acquired on a Siemens Magnetom Prisma 3 Tesla MRI scanner (Siemens, Erlangen, German) at the Baker Heart and Diabetes Institute, Melbourne, Australia, at 6 -11 weeks following injury. DWI data and one structural image (T1w) were acquired in a single session, using a 64-channel head coil. Structural T1-weighted images were acquired using the magnetisation prepared rapid gradient-echo (MPRAGE) sequence (240 x 256 mm acquisition matrix; FOV = 256mm; 176 contiguous slices; 1mm^3^ isotropic voxel size; TR/TE = 2300/2.96 ms).

DWI data were acquired in the anterior-posterior phase encoded direction, with multiband accelerated EPI sequences for multishell acquisition (128 x 128 mm acquisition matrix; FOV = 256mm; 75 contiguous slices; 64 volumes b = 1000 s/mm^2^, 64 volumes b = 3000 s/mm^2^, 4 volumes b = 0 s/mm^2^; 2mm^3^ isotropic resolution; TR/TE = 4800/88 ms; MB factor = 3). Two mTBI and 1 TC participants were removed from analysis due to an error in the acquisition protocol for the DWI sequences. All scans were reviewed by an experienced neuroradiologist for potential incidental findings. A brain tumour was identified in one mTBI participant, who was subsequently excluded from analysis.

### MRI quality control and preprocessing

All structural images were visually inspected for any potential artifacts including movement. Each subject’s T1 images were pre-processed using the standard *recon-all* processing pipeline, in the FreeSurfer analysis suite release 7.4.1 (Fischl, 2012; https://surfer.nmr.mgh.harvard.edu/). This pipeline performs non-parametric non-uniform intensity normalisation and skull stripping. Estimated intracranial volume (ICV) was calculated for each participant using the total Intracranial Volume output from *recon-all.* All structural reconstructions underwent manual quality control according to the method suggested by Klapwijk et al. (2019). All structural reconstructions satisfied quality control, and no manual editing of surface boundaries was undertaken.

DWI data were pre-processed using an established pipeline which uses a combination of the MRtrix3 software package (version 3.0.4; Brain Research Institute, Melbourne, Australia, www.mrtrix.org) and FSL (FMRIB’s Software Library; https://fsl.fmrib.ox.ac.uk/fsl). Data were denoised and corrected for Gibbs ringing (Kellner et al., 2016; Veraart et al., 2016). Susceptibility- induced geometric distortions were corrected using Synb0-DisCo (Schilling et al., 2019; 2020), and bias field was corrected using the N4 algorithm (Tustison et al., 2010). We manually checked images following preprocessing to ensure there were no susceptibility induced or motion distortions.

### Thalamus segmentation

Lateralised whole and regional thalamic segmentations were estimated using a Bayesian joint structural and DWI data segmentation method, which combines information from a previous probabilistic atlas incorporating histologically identified nuclei together with likelihood models for both structural and diffusion data (Tregidgo et al., 2023). The algorithm requires the bias field corrected image and whole brain parcellation yielded by the *recon-all* pipeline, and a diffusion tensor imaging (DTI) model as fitted by TRACULA (Maffei et al, 2021; Yendiki et al., 2011). The DTI model was fitted to the b = 1000 s/mm^2^ shell. All segmentations were visually inspected by M.B and J.Y; the latter is a trained neurosurgeon with over 10 year’s experience in neuroimaging research. One TC participant was removed due to clear false positive segmentation into the right internal capsule.

To avoid over-inflation of statistical estimates for smaller subnuclei volumes, the 24 nuclei labels from each hemisphere were merged to create subregions. Label merging decisions were informed by the histological atlas utilised in the development of the segmentation method (Morel et al., 1997; Tregidgo et al., 2023). The thalamic subgroups utilised are also broadly consistent with commonly employed grouping based on anatomical location (Grodd et al., 2020). Figure 1 presents an example segmentation for one participant, along with the labels merged to create thalamic subregions. Volumes were extracted for each subnuclei and summed according to labels to yield nuclei regions. Estimations for all segmented subnuclei were summed to produce lateralised whole thalamus volumes.

**Figure 1.**
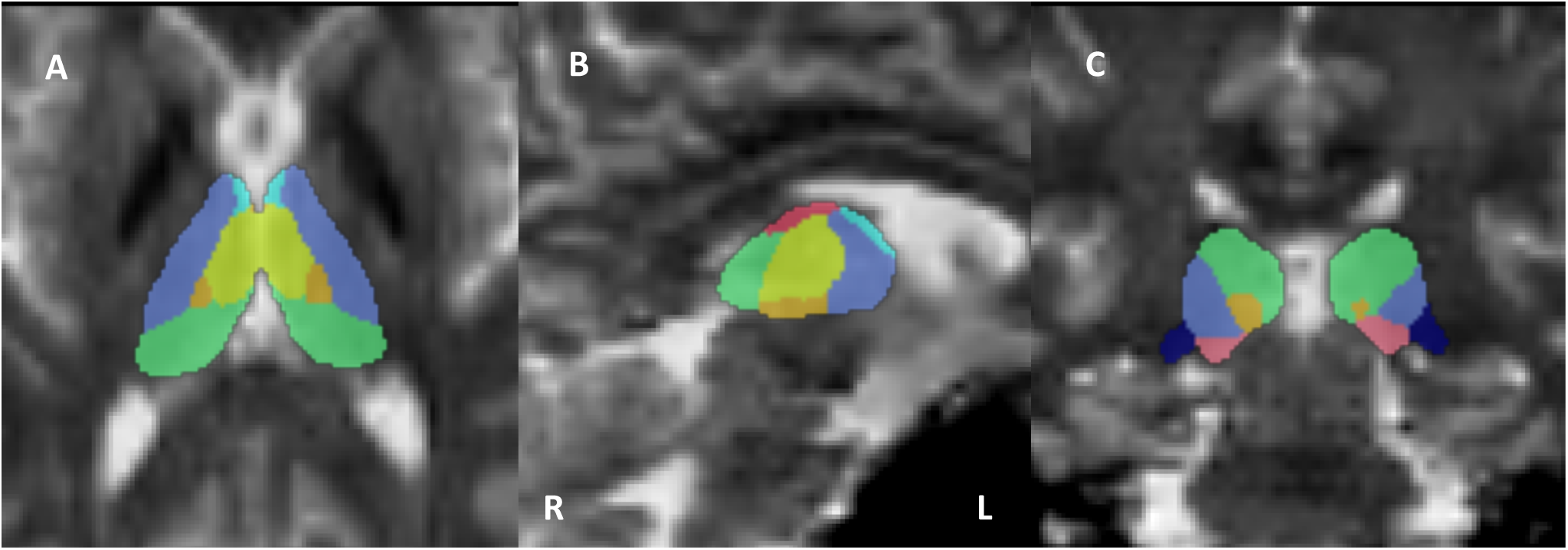
Example segmentation and nuclei label merging overlayed on diffusion tensor fractional anisotropy map

**Fig. 1.**
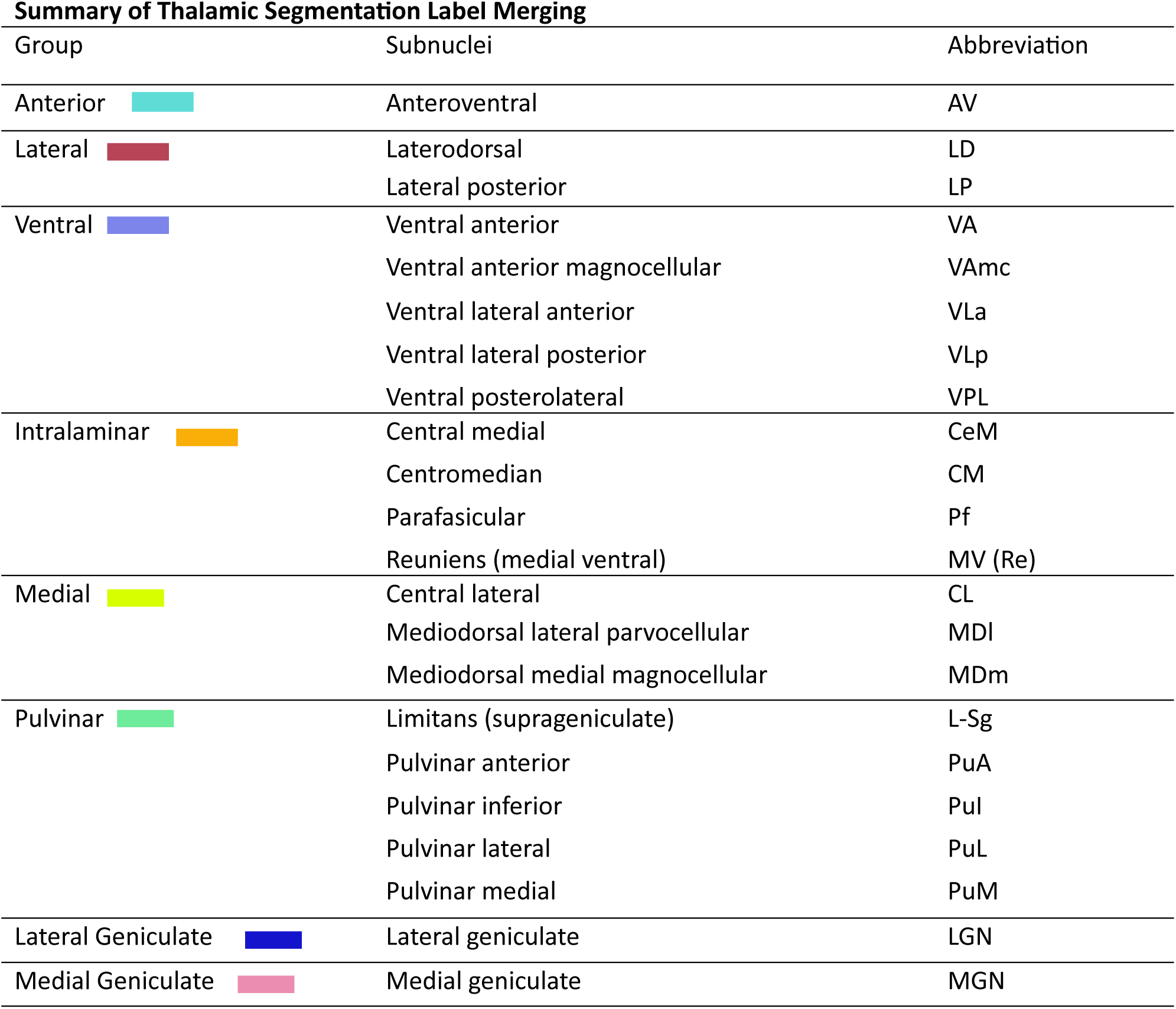
Axial (a), sagittal (b) and coronal (c) views of an example thalamus segmentation obtained from a participant. Anterior region (light blue); medial region (yellow); lateral region (dark pink); intralaminar (orange); ventral region (purple); lateral geniculate nucleus (dark blue); medial geniculate nucleus (light pink).

### Supplementary analysis: NODDI

In order to assist interpretation of findings of the volume analyses, a NODDI model was computed in the same spatial location as the extracted thalamic nuclei volumes. Pre-processed diffusion images were up-sampled to 0.7mm^3^ using nearest neighbour interpolation prior to fitting the NODDI model. The NODDI Matlab Toolbox (Version 1.0.5) was used to fit the multishell DWI data to the NODDI tissue model. This toolbox computes parametric maps of the three NODDI metrics: ODI, ISO, and NDI (Zhang et al., 2012). In its original form, the NODDI default model parameters are optimised for white matter. Since parallel diffusivity varies between white and grey matter, we applied an intrinsic parallel diffusivity value of (1.1 × 10^−3^ mm^2^/s), as this is in the range of values shown to be more optimal for grey matter structures (Fukutomi et al., 2018; Guerrero et al., 2019). Following model computation, average subregion parameters were then estimated from binarized subgroup masks derived from segmentations.

### Statistical analyses

The frequentist approach to hypothesis testing provides a limited measure of the strength of evidence against the null hypothesis (of no group difference) and typically relies on large sample sizes to attain meaningful results (Makowski et al., 2019). To overcome these limitations, a Bayesian analysis framework was adopted for all statistical analyses. All analyses were conducted in JASP (version 0.18.2; Jasp Team, 2023) and R (version 3.3.0). Where appropriate, we report the median of the posterior distribution and its associated credible interval (95% CI), along with the Bayes Factors (BF). Group differences in demographic and injury-related variables were evaluated using Bayesian Independent Samples T-tests and Bayesian Tests of Association (sex). For all analyses, BF_10_ values greater than 3 were considered as providing moderate and increasing evidence in favour of the alternative hypothesis; BF_10_ values less than 1/3 were considered as moderate and increasing evidence in support of the null model (i.e. strong evidence of no difference between groups); BF_10_ values between 1/3 and 3 were considered as offering inconclusive support for or against the null hypothesis (Makowski et al., 2019). There were no missing data for any demographic or injury-related variables. Univariate outliers were classified as scores > 3.29 standard deviations above or below the mean (Tabachnick & Fidell, 2014). No univariate outliers were identified for any of the variables. The assumptions of normality and homogeneity of variance were met.

To determine the inclusion of covariates in the main analyses, Bayesian correlations were performed to evaluate the association between lateralised thalamus subregion volumes and age, and ICV, and a Bayesian Independent Samples T-Test was performed to evaluate the association between sex and subregion volumes. All covariate analyses were performed within each group separately. As reported in the Supplementary material Table 1, in both groups there was evidence for a strong relationship between age and volume, and ICV and volume across multiple subregions. There was also evidence of a relationship in both groups between sex and thalamus volumes across multiple regions. To evaluate the relationship between time since injury and thalamus volumes, we performed Bayesian correlations between lateralised subregion volumes and days since injury. As reported in the Supplementary material Table 1, within each group there was evidence for no relationship between time since injury and volume for many of the thalamic subregions (BF_10_ < 1/3), and inconclusive evidence for the remaining subregions (1/3 < BF_10_ ≥ 3). Therefore, age, sex and ICV were included as covariates in subsequent analyses to control for any influence on thalamus subregion volumes. For consistency across models, the same covariates were included in the models examining group differences in NODDI metrics.

To examine whether there were differences in regional thalamus volumes and NODDI metrics between mTBI and TC groups, a series of Bayesian linear regression models were fitted and the BF_10_ for the alternative hypothesis of a group difference in the outcome variable were computed. Priors over parameters were all set as normal, and model priors were uniform. Fixed effects were scaled to 0.5 and covariates to 0.354, as per the JASP defaults. For all volumetric and NODDI analyses, the null model included sex, age, and ICV as predictors, and the alternative model contained these predictors plus diagnostic group.

## Results

### Demographic and injury-related variables

The final dataset included 67 participants (mTBI, n = 39) who completed the protocol at 6 - 11 weeks following injury (range: 40 – 78; mean: 57 days; sd: 11). The groups were well-matched in terms of demographic and injury-related characteristics. As presented in Table 1, there was no evidence in support of a difference in age, level of education, ratio of female to male biological sex, accident type, or time elapsed between injury and assessment between the mTBI and TC groups.

**Table 1.**
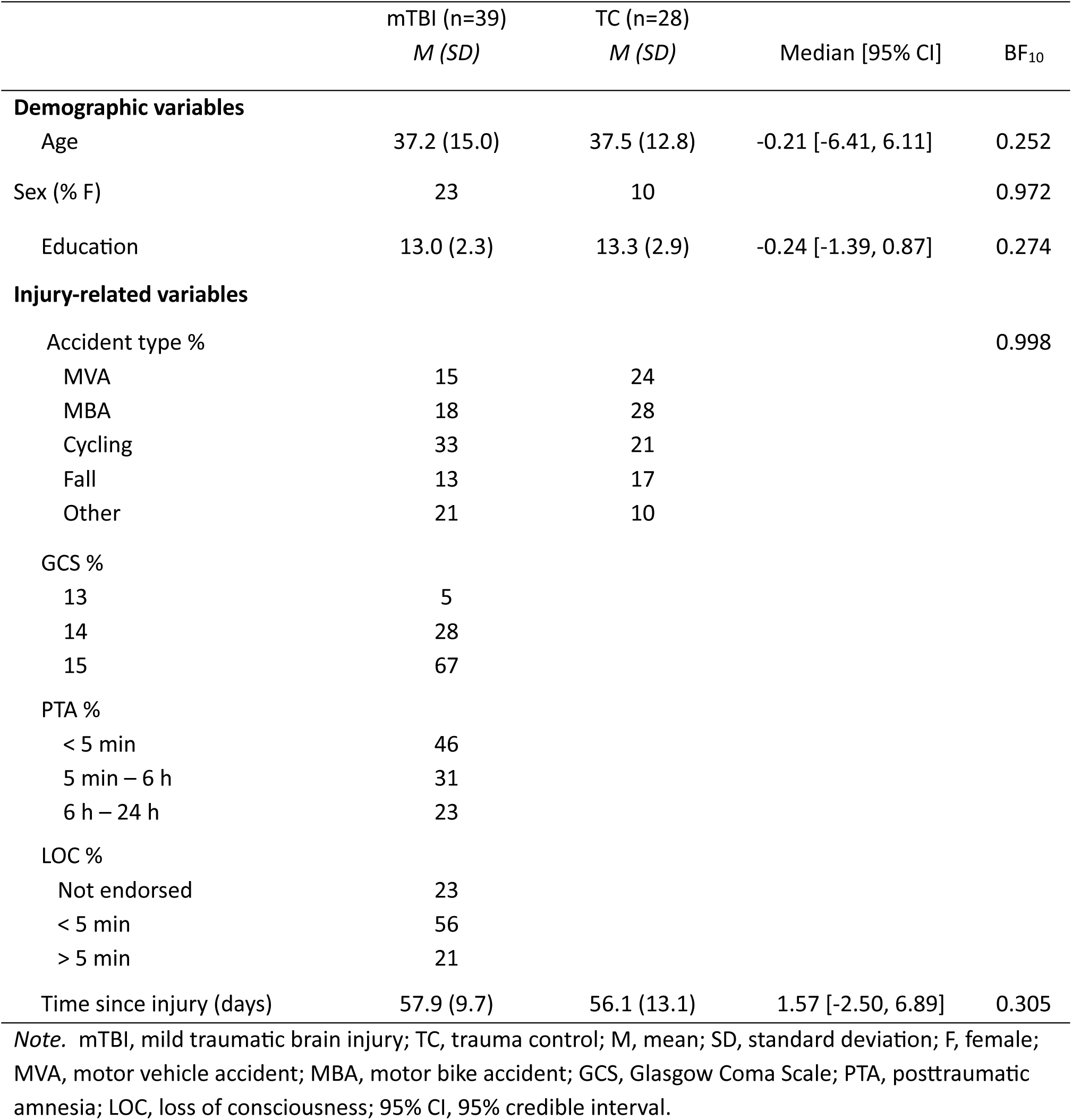
Demographic and injury-related variables for mTBI and TC groups.

|  | mTBI (n=39)<br>*M (SD)* | TC (n=28)<br>*M (SD)* | Median [95% CI] | BF<sub>10</sub> |
| --- | --- | --- | --- | --- |
| **Demographic variables** |  |  |  |  |
| Age | 37.2 (15.0) | 37.5 (12.8) | -0.21 [-6.41, 6.11] | 0.252 |
| Sex (% F) | 23 | 10 |  | 0.972 |
| Education | 13.0 (2.3) | 13.3 (2.9) | -0.24 [-1.39, 0.87] | 0.274 |
| **Injury-related variables** |  |  |  |  |
| Accident type % |  |  |  | 0.998 |
| MVA | 15 | 24 |  |  |
| MBA | 18 | 28 |  |  |
| Cycling | 33 | 21 |  |  |
| Fall | 13 | 17 |  |  |
| Other | 21 | 10 |  |  |
| GCS % |  |  |  |  |
| 13 | 5 |  |  |  |
| 14 | 28 |  |  |  |
| 15 | 67 |  |  |  |
| PTA % |  |  |  |  |
| < 5 min | 46 |  |  |  |
| 5 min – 6 h | 31 |  |  |  |
| 6 h – 24 h | 23 |  |  |  |
| LOC % |  |  |  |  |
| Not endorsed | 23 |  |  |  |
| < 5 min | 56 |  |  |  |
| > 5 min | 21 |  |  |  |
| Time since injury (days) | 57.9 (9.7) | 56.1 (13.1) | 1.57 [-2.50, 6.89] | 0.305 |

*Note.* mTBI, mild traumatic brain injury; TC, trauma control; M, mean; SD, standard deviation; F, female; MVA, motor vehicle accident; MBA, motor bike accident; GCS, Glasgow Coma Scale; PTA, posttraumatic amnesia; LOC, loss of consciousness; 95% CI, 95% credible interval.

### Differences in thalamic subfield volume between mTBI and TC

Distributions of thalamus subregion volumes are presented in Figure 2. Results from the Bayesian linear models for group status predicting subregion volume are presented in Table 2.

**Figure 2.**
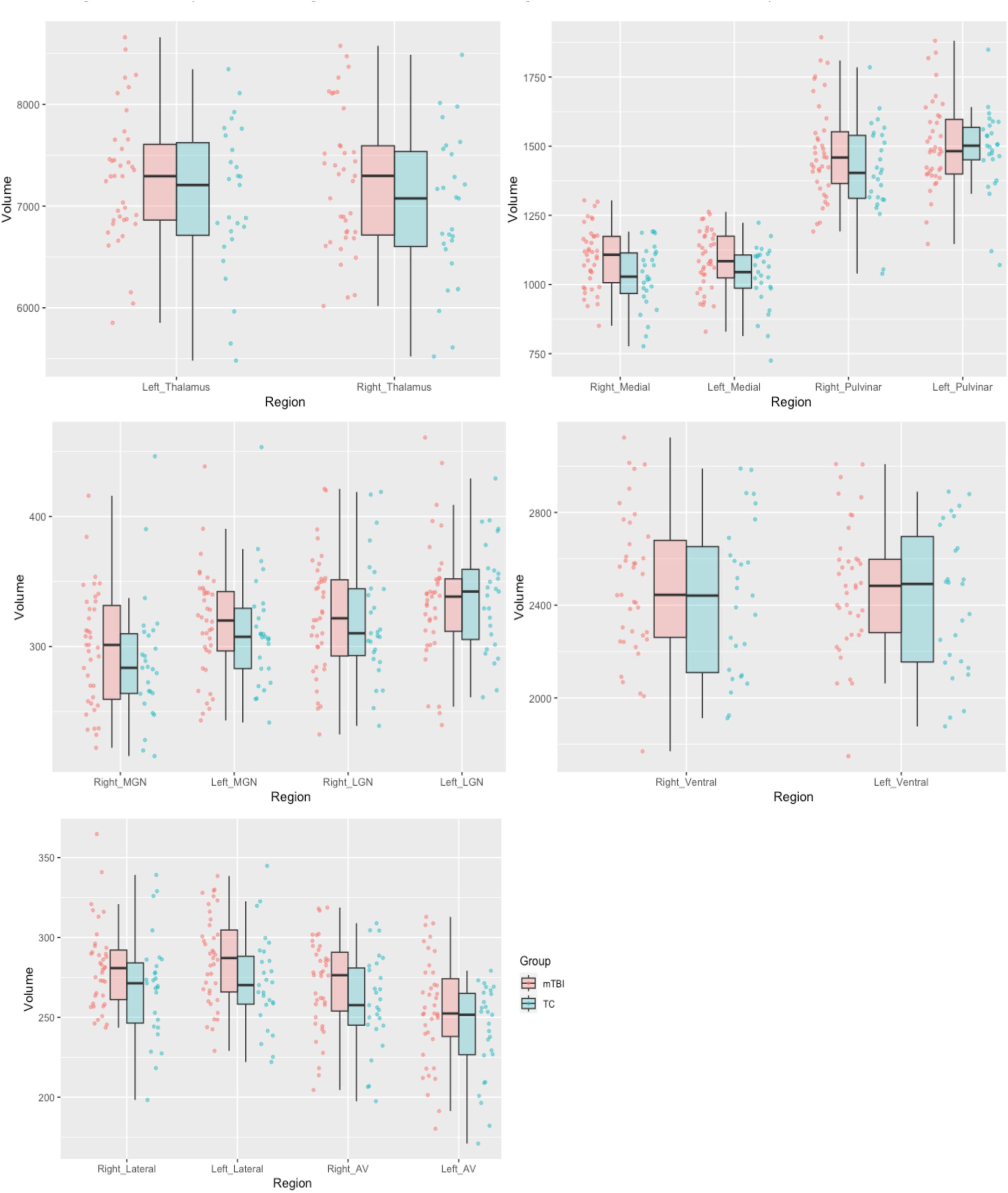
Box plots showing distributions of subregion volumes in mTBI compared to TCs *Note.* LGN, lateral geniculate nucleus; MGN, medial geniculate nucleus; mTBI, mild traumatic brain injury; TC, trauma control.

**Table 2.**
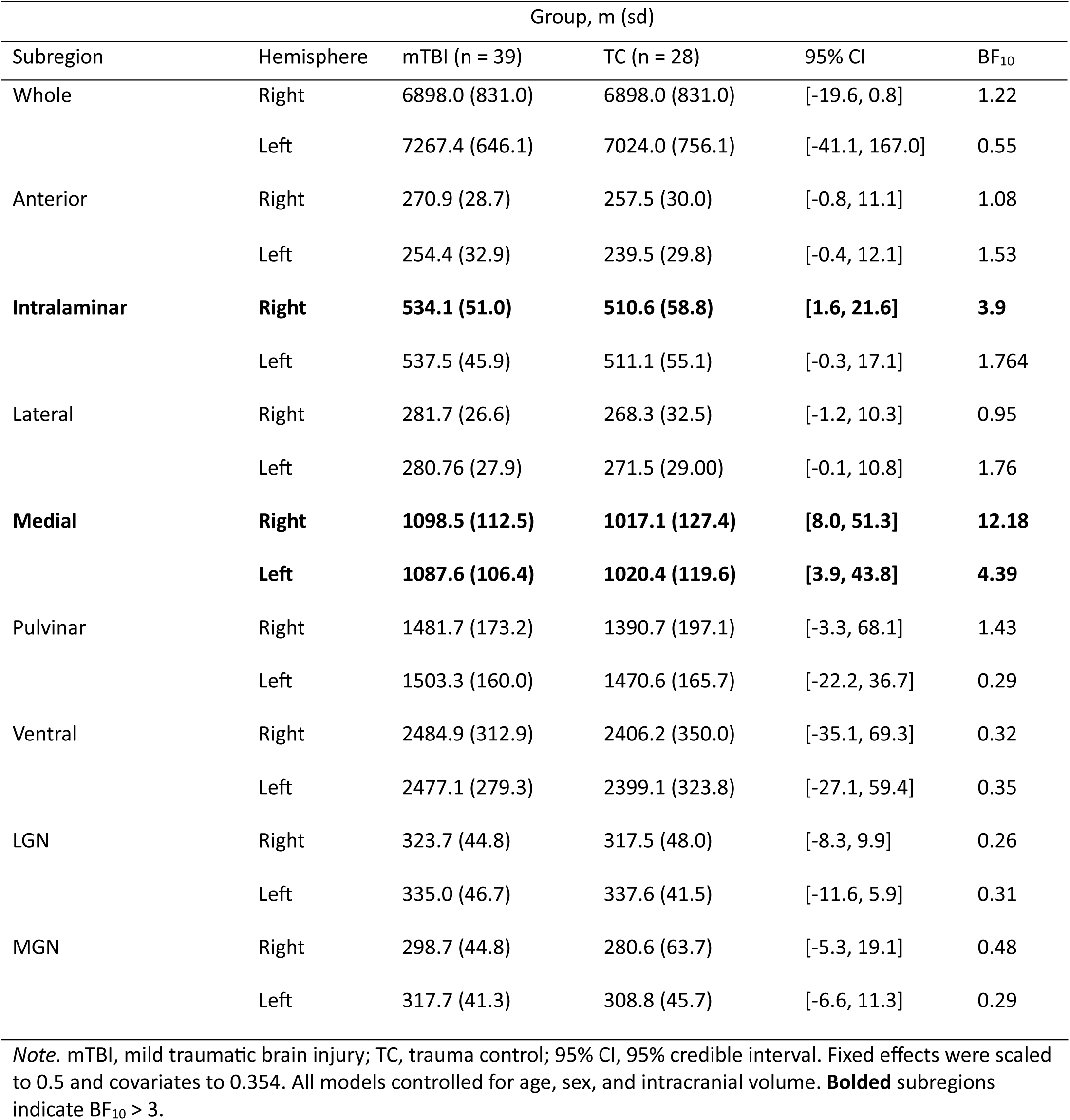
Linear models for group status predicting thalamic subregion volume.

| Subregion | Hemisphere | Group, m (sd) |  | 95% CI | BF<sub>10</sub> |
| --- | --- | --- | --- | --- | --- |
|  |  | mTBI (n = 39) | TC (n = 28) |  |  |
| Whole | Right | 6898.0 (831.0) | 6898.0 (831.0) | [-19.6, 0.8] | 1.22 |
|  | Left | 7267.4 (646.1) | 7024.0 (756.1) | [-41.1, 167.0] | 0.55 |
| Anterior | Right | 270.9 (28.7) | 257.5 (30.0) | [-0.8, 11.1] | 1.08 |
|  | Left | 254.4 (32.9) | 239.5 (29.8) | [-0.4, 12.1] | 1.53 |
| **Intralaminar** | **Right** | **534.1 (51.0)** | **510.6 (58.8)** | **[1.6, 21.6]** | **3.9** |
|  | Left | 537.5 (45.9) | 511.1 (55.1) | [-0.3, 17.1] | 1.764 |
| Lateral | Right | 281.7 (26.6) | 268.3 (32.5) | [-1.2, 10.3] | 0.95 |
|  | Left | 280.76 (27.9) | 271.5 (29.00) | [-0.1, 10.8] | 1.76 |
| **Medial** | **Right** | **1098.5 (112.5)** | **1017.1 (127.4)** | **[8.0, 51.3]** | **12.18** |
|  | **Left** | **1087.6 (106.4)** | **1020.4 (119.6)** | **[3.9, 43.8]** | **4.39** |
| Pulvinar | Right | 1481.7 (173.2) | 1390.7 (197.1) | [-3.3, 68.1] | 1.43 |
|  | Left | 1503.3 (160.0) | 1470.6 (165.7) | [-22.2, 36.7] | 0.29 |
| Ventral | Right | 2484.9 (312.9) | 2406.2 (350.0) | [-35.1, 69.3] | 0.32 |
|  | Left | 2477.1 (279.3) | 2399.1 (323.8) | [-27.1, 59.4] | 0.35 |
| LGN | Right | 323.7 (44.8) | 317.5 (48.0) | [-8.3, 9.9] | 0.26 |
|  | Left | 335.0 (46.7) | 337.6 (41.5) | [-11.6, 5.9] | 0.31 |
| MGN | Right | 298.7 (44.8) | 280.6 (63.7) | [-5.3, 19.1] | 0.48 |
|  | Left | 317.7 (41.3) | 308.8 (45.7) | [-6.6, 11.3] | 0.29 |

*Note.* mTBI, mild traumatic brain injury; TC, trauma control; 95% CI, 95% credible interval. Fixed effects were scaled to 0.5 and covariates to 0.354. All models controlled for age, sex, and intracranial volume. **Bolded** subregions indicate BF<sub>10</sub> > 3.

Although there was no difference between the mTBI and TC groups in ICV, there was evidence for increased volume in the mTBI group compared to TCs specifically for the bilateral medial thalamus and right intralaminar thalamus regions (BF_10_ > 3). There was evidence for no group differences in volume in the right ventral, left pulvinar, bilateral LGN and left MGN subregions (BF_10_ < 1/3). For the remaining subregions, the evidence was inconclusive, with no evidence for or against the null/alternative hypotheses provided (1/3 < BF_10_ ≥ 3).

### Differences in NODDI metrics between mTBI and TC

A NODDI model was computed to help inform interpretation of the volume increases noted above. Results from the Bayesian linear models for group status predicting NDI, ODI and ISO are presented in Table 3. Of the subregions which showed increased volume, there was no evidence to support group differences in NODDI metrics. There was good evidence to support no group differences for ODI, NDI, and ISO in the right medial thalamus, and for NDI and ISO in the left medial thalamus. There was inconclusive evidence to support either the null or alternative hypothesis for group differences in ODI in the left medial thalamus, and for ODI, NDI, and ISO in the right intralaminar subregion.

**Table 3.** Group differences in Thalamic subnuclei volume: output from Bayesian linear regression models for group status predicting NODDI metrics, controlling for sex, age, and estimated intracranial volume (reference group = mTBI)

| Subregion | Hemisphere | ISO | NDI | ODI |
| --- | --- | --- | --- | --- |
|  |  | BF<sub>10</sub> | BF<sub>10</sub> | BF<sub>10</sub> |
| Whole | Right | 0.26 | 1.18 | 0.27 |
|  | Left | 0.26 | 0.31 | 0.36 |
| Anterior | Right | 0.44 | 0.51 | 0.35 |
|  | Left | 0.30 | 0.34 | 0.44 |
| Intralaminar | Right | 0.43 | 1.03 | 0.39 |
|  | Left | 0.83 | 0.77 | 0.33 |
| Lateral | Right | 0.26 | 0.27 | 0.25 |
|  | Left | 0.27 | 0.28 | 0.34 |
| Medial | Right | 0.28 | 0.27 | 0.30 |
|  | Left | 0.28 | 0.25 | 0.40 |
| Pulvinar | Right | 0.29 | 0.51 | 0.38 |
|  | Left | 0.26 | 0.25 | 0.27 |
| Ventral | Right | 0.26 | 0.68 | 0.43 |
|  | Left | 0.26 | 0.27 | 0.32 |
| LGN | Right | 0.32 | 1.34 | 0.25 |
|  | Left | 0.26 | 0.40 | 0.28 |
| MGN | Right | 0.32 | 0.99 | 0.39 |
|  | Left | 0.26 | 0.64 | 0.44 |

*Note.* NDI, neurite density index; ISO, volume of isotropic free water component; ODI, orientation dispersion index; 95% CI, 95% credible interval. Fixed effects were scaled to 0.5 and covariates to 0.354. All models controlled for age, sex, and intracranial volume.

## Discussion

This study aimed to investigate thalamic structure following mTBI. We used a multimodal imaging approach to segment the thalamus into its constituent subnuclei, then extracted subregion volumes, and as a supplementary analysis computed microstructural NODDI parameters for the same regions. Our findings show that at approximately 8 weeks following injury, mTBI is associated with greater volume in specific thalamic subregions, which suggests thalamic structural abnormalities following mTBI persist beyond the acute period following injury.

Specifically, we observed that mTBI is associated with increased grey matter volume compared to TCs in the bilateral medial thalamus and the right intralaminar nuclei subregions. The strongest evidence for group differences in thalamic subregion volume was observed in the bilateral medial thalamus region. To the best of our knowledge, only one prior volumetric study has examined thalamic subnuclei volume following mTBI (Woodrow et al., 2023). This study showed no differences in subregion volumes between mTBI and controls at an average of 2-weeks post injury, although some participants underwent imaging as early as 72 hours following the mTBI event. Our finding of increased medial thalamus volume at an average of 2 months following injury, in addition to longitudinal work showing changes in whole thalamus volume occur over the months post-injury, suggests volume changes may occur over a protracted period rather than as an immediate acute response to injury (Munivenkatappa et al., 2016; Zagorchev et al., 2016; Zhou et al., 2021).

Specifically, one prior study reported an increase in whole thalamus volume over time following mTBI, and greater volume in the mTBI group compared to controls at 6 – 7 months following injury (Munivenkatappa et al., 2016). Increased volume following mTBI has also been demonstrated in other brain structures, including the ventromedial prefrontal cortex, right fusiform cortex, dorsal anterior cingulate cortex and dorsal posterior cingulate cortex (Killgore et al., 2016; Niu et al., 2020).

This study demonstrates differential sensitivity of specific thalamic subnuclei to the effects of mTBI. These selective effects could be due to differences in cytoarchitecture and myeloarchitecture, which have been shown to influence susceptibility to injury in mTBI (Taber & Hurley, 2013). The pattern of findings demonstrates the importance of considering structural change within subregions of the thalamus, particularly considering the variable contribution of different subregions to function. For example, the medial thalamus, defined in this study as the central lateral nuclei and the lateral and medial regions of the mediodorsal nuclei, is a higher-order structure which receives few or no sensory inputs (Mitchell & Charkraborty, 2013). Robust bi-directional connections with the prefrontal cortex enable its critical role in cognitive functions including working memory, attention control, set-shifting, and decision making (Georgescu et al., 2020; Parnaudeau et al., 2018). Given its contribution to cognitive functions, structural change in the medial thalamus could contribute to poorer cognitive outcome subacute period (Hacker et al., 2023). Future work should investigate this by examining associations between subnuclei structural metrics and relevant outcome measures in mTBI samples.

Increases in volume identified using MRI-based methods are non-specific and can reflect contributions of multiple components including neurogenic inflammation, increased presence of astrocytes, glial cell genesis, excess myelin debris, and excessive remyelination (Cole et al., 2016). In an effort to clarify the possible biological mechanism underpinning the increased volume observed here, we also conducted an exploratory NODDI analysis. Of the subregions which showed greater volume associated with mTBI, there was evidence for no differences in NODDI metrics between groups for all right medial NODDI metrics, and for all left medial NODDI metrics aside from ODI (BF_10_ < 1/3). There was inconclusive evidence for group differences in all NODDI metrics for the right intralaminar region, and for the left medial ODI metric (1/3 < BF_10_ ≥ 3). While the NODDI model has been shown to relate to cellular properties including oedema and atrophy, the metrics are non- exhaustive in terms of microstructural tissue features they are assumed to reflect. Therefore, it is likely that the mechanisms underpinning the increased volume observed here are not well captured by this model. For example, myelin-related changes would be poorly captured by the NODDI model, as the echo time for the DWI sequences used in this study are too long to capture the rapidly decaying diffusion signal from the myelin water compartment which means myelin-specific information from neural tissue cannot be attained (Ma et al., 2020). It is plausible the increased volume observed here could be attributed to excessive remyelination or myelin debris yet to be cleared. Distal to sites of axonal disconnection, myelin sheaths can collapse and propagate neuroinflammation due to a protracted clearance of debris and subsequent microglial activation (Armstrong et al., 2016; Yang et al., 2011). Indeed, increased thalamic myelin content has been shown in mTBI groups 3-months following injury, which has been suggested to reflect axon neuropathology or disordered remyelination leading to excessive myelination (Spider et al., 2018). It would be beneficial for future research to include imaging sequences that can be modelled to infer myelin properties, so thalamic volumetrics and myelination can be examined within the same sample of participants.

The current study presents some possible limitations. Firstly, while the statistical analyses utilised are more robust to small sample sizes compared to traditional frequentist methods, the modest participant sample size in this study may be underpowered to detect changes in thalamic structure. Additionally, the NODDI findings presented here should be noted with caution. NODDI relies on simplified assumptions about the tissue properties it attempts to probe, which are present in the model as fixed parameters (Kamiya et al., 2020). Some previous work has suggested the variation of these parameters influences the model fit and the extracted metrics (Guerroro et al.,2019). While we opted for an intrinsic diffusivity value utilised elsewhere, given the paucity of research applying the NODDI model to grey matter structures, this value has not been widely validated.

There are some notable strengths of the present study. This is the first application of a joint structural and diffusion MRI thalamic segmentation algorithm within an mTBI sample (Tregigdo et al., 2023). Many widely employed thalamic segmentation methods warp histologically-derived atlases to the T1 images alone, which relies on negligible contrast differences within the thalamus to define boundaries between subnuclei. The addition of diffusion data provides complementary information to improve subnuclei boundary placement. Given the robust method employed here is freely available as part of the FreeSurfer imaging analysis suite, it is hoped future studies will employ the segmentation algorithm used here. The recruitment of participants who were premorbidly healthy is also an important methodological strength in the present study, as it isolates the observed increased volume to the effects of the mTBI.

## Conclusions

For the first time, this study reports structural abnormalities in the bilateral medial thalamus and right intralaminar subregions at approximately 8-weeks following an mTBI event. This suggests structural recovery is not yet complete beyond the acute period following injury. Importantly, the results show there is differential sensitivity between thalamic nuclei to the effects of mTBI, which highlights the importance of addressing the constituent nuclei of the thalamus in neuroimaging investigations that attempt to probe injury-related neuropathology. While we computed a NODDI model to help interpret these results, our findings suggest the biological underpinnings of the increased volume are not well captured by the NODDI metrics and are more accurately explained by other neuropathological mechanisms. Putative mechanisms underpinning the increased volume are disordered remyelination, or myelin debris yet to be cleared. Future studies should investigate this by utilising neuroimaging methods which are more sensitive to myelin-specific information, such as the T1w/T2w ratio maps (Dipnall et al., 2024). It is also important future work extends the findings presented here by determining whether structural changes in thalamic subregions have specific effects on mTBI outcomes.

## Acknowledgements

The authors thank student researchers who assisted in data collection, the Baker Heart and Diabetes Institute imaging group, Jian Chen for technical support, and those who participated in this study. The research and analyses were conducted within the Developmental Imaging research group, Murdoch Children’s Research Institute, supported by the Victorian Government’s Operational Infrastructure Support Program, and The Royal Children’s Hospital Foundation devoted to raising funds for research at The Royal Children’s Hospital.

## Author Contributions

All authors had full access to all the data in the study and take responsibility for the integrity of the data and the accuracy of the data analysis. *Conceptualisation,* M.B, R.B, J.Y, M.S, J.A; *Methodology,* M.B, R.B, J.Y, M.S, J.A; *Investigation,* M.B, J.Y, R.B; *Software,* M.B, R.B; *Formal analysis,* M.B, *Writing – Original draft,* M.B; *Writing- Reviewing & Editing,* R.B, J.Y, M.S, J.A; *Funding acquisition,* J.A.

## Abbreviations

DWI: Diffusion Weighted Imaging
MRI: Magnetic Resonance Imaging
MPRAGE: Magnetisation Prepared Rapid Gradient-Echo
mTBI: Mild Traumatic Brain Injury
NODDI: Neurite Orientation and Dispersion Density Imaging
TC: trauma control

## Supplementary Material

**Supplementary Table 1.** Results of Bayesian Correlations between thalamus subregion volumes and age, ICV, and days since injury.

|  |  | Age |  |  |  | ICV |  |  |  | Days since injury |  |  |  |
| --- | --- | --- | --- | --- | --- | --- | --- | --- | --- | --- | --- | --- | --- |
|  |  | mTBI |  | TC |  | mTBI |  | TC |  | mTBI |  | TC |  |
| Subregion | Hemisphere | *r* | BF<sub>10</sub> | *r* | BF<sub>10</sub> | *r* | BF<sub>10</sub> | *r* | BF<sub>10</sub> | *r* | BF<sub>10</sub> | *r* | BF<sub>10</sub> |
| Anterior | Right | -0.53, | 58.2 | -0.32 | 0.86 | 0.19 | 0.40 | 0.33 | 0.97 | -0.25 | 0.61 | -0.14 | 0.30 |
|  | Left | -0.57 | 212.1 | -0.49 | 6.43 | 0.13 | 0.27 | 0.22 | 0.43 | -0.18 | 0.36 | -0.19 | 0.37 |
| Intralaminar | Right | -0.22 | 0.47 | -0.26 | 0.56 | 0.62 | 963.29 | 0.56 | 21.28 | -0.30 | 1.05 | 0.21 | 0.41 |
|  | Left | -0.23 | 0.52 | -0.16 | 0.32 | 0.63 | 1597.14 | 0.64 | 140.42 | -0.26 | 0.68 | 0.13 | 0.29 |
| Lateral | Right | 0.13 | 0.27 | 0.15 | 0.31 | 0.48 | 18.81 | 0.72 | 14.2e+3 | -0.06 | 0.20 | 0.02 | 0.24 |
|  | Left | 0.07 | 0.22 | 0.14 | 0.30 | 0.56 | 137.23 | 0.71 | 10.8e+3 | -1.2e-4 | 0.21 | -0.04 | 0.24 |
| Medial | Right | -0.24 | 0.57 | -0.14 | 0.30 | 0.58 | 281.71 | 0.67 | 297.07 | -0.16 | 0.31 | 0.03 | 0.24 |
|  | Left | -0.29 | 0.90 | -0.02 | 0.24 | 0.60 | 450.77 | 0.69 | 630.24 | -0.20 | 0.42 | 0.05 | 0.24 |
| Pulvinar | Right | -0.30 | 0.99 | -0.49 | 6.57 | 0.5 | 33.66 | 0.34 | 1.07 | -0.16 | 0.32 | 0.30 | 0.73 |
|  | Left | -0.29 | 0.98 | -0.32 | 0.83 | 0.56 | 134.78 | 0.49 | 6.47 | -0.15 | 0.30 | 0.26 | 0.56 |
| Ventral | Right | 0.30 | 1.03 | 0.20 | 0.38 | 0.71 | 44.8e+6 | 0.78 | 22.9e+4 | -0.10 | 0.24 | 0.13 | 0.29 |
|  | Left | 0.22 | 0.48 | 0.22 | 0.43 | 0.76 | 61.1e+7 | 0.84 | 45.1e+5 | -0.03 | 0.20 | 0.06 | 0.25 |
| LGN | Right | 0.20 | 0.40 | 0.10 | 0.27 | 0.47 | 15.50 | 0.61 | 62.16 | -0.12 | 0.26 | 0.23 | 0.45 |
|  | Left | 0.09 | 0.22 | 0.24 | 0.48 | 0.43 | 6.87 | 0.69 | 552.60 | -0.04 | 0.21 | 0.20 | 0.38 |
| MGN | Right | 0.03 | 0.40 | -0.30 | 0.71 | 0.46 | 13.30 | 0.15 | 0.31 | -0.19 | 0.39 | 0.33 | 0.96 |
|  | Left | 0.20 | 0.41 | -0.02 | 0.24 | 0.52 | 56.31 | 0.46 | 4.01 | -0.02 | 0.20 | 0.26 | 0.56 |

*Note.* mTBI, mild traumatic brain injury; TC, trauma control; 95% CI, 95% credible interval; LGN, lateral geniculate nucleus; MGN, medial geniculate nucleus. Fixed effects were scaled to 0.5

**Supplementary Table 2.** Results of Bayesian Independent Samples T-Test for association between sex and thalamus subregion volume (reference group = male)

| Subregion | Hemisphere | mTBI |  | TC |  |
| --- | --- | --- | --- | --- | --- |
|  |  | 95% CI | BF<sub>10</sub> | 95% CI | BF<sub>10</sub> |
| Anterior | Right | [-18.26, 27.56] | 0.52 | [-22.78, 8.03] | 0.72 |
|  | Left | [-8.33, 14.23] | 0.40 | [-23.45, 7.47] | 0.78 |
| Intralaminar | Right | [-4.24, 31.96] | 1.15 | [33.13, 24.80] | 0.50 |
|  | Left | [-5.08, 26.87] | 0.88 | [32.68, 21.55] | 0.51 |
| Lateral | Right | [0.05, 18.89] | 2.85 | [12.26, 19.41] | 0.52 |
|  | Left | [2.83, 22.71] | 10.75 | [-9.02, 20.63] | 0.62 |
| Medial | Right | [-4.83, 75.13] | 1.75 | [63.21, 61.05] | 0.48 |
|  | Left | [-7.78, 67.37] | 1.33 | [-54.08, 61.57] | 0.48 |
| Pulvinar | Right | [17.85, 103.70] | 0.95 | [122.79, 72.07] | 0.53 |
|  | Left | [-4.26, 110.20] | 2.22 | [78.47, 84.22] | 0.48 |
| Ventral | Right | [46.17, 262.95] | 23.36 | [-77.73, 296.53] | 0.88 |
|  | Left | [36.94, 232.40] | 18.54 | [-81.96, 261.91] | 0.78 |
| LGN | Right | [-6.01, 24.98] | 0.76 | [-20.51, 26.95] | 0.50 |
|  | Left | [-6.07, 25.65] | 0.78 | [16.45, 25.58] | 0.52 |
| MGN | Right | [-8.74, 22.05] | 0.51 | [-37.41, 25.78] | 0.51 |
|  | Left | [-4.03, 24.98] | 1.03 | [18.26, 27.56] | 0.52 |

*Note.* mTBI, mild traumatic brain injury; TC, trauma control; 95% CI, 95% credible interval; LGN, lateral geniculate nucleus; MGN, medial geniculate nucleus. Fixed effects were scaled to 0.5

